# Whole Proteome Clustering of 2,307 Genomes Reveals Remarkable Conservation of Four Proteins Among Proteobacteria While Revealing Significant Annotation Issues

**DOI:** 10.1101/352856

**Authors:** Svetlana Lockwood, Kelly A. Brayton, Jeff A. Daily, Shira L. Broschat

**Author notes:** Corresponding author (SL).

## Abstract

To explore the concept of a minimal gene set, we clustered 8.76 M protein sequences deduced from 2,307 completely sequenced Proteobacterial genomes. To our knowledge this is the first study of this scale. Clustering resulted in 707,311 clusters of which 224,442 ranged in size from 2 to 2,894 sequences. The resulting clusters allowed us to ask the question: Is a set of proteins conserved across all Proteobacteria? We chose four essential proteins, the chaperonin GroEL, DNA dependent RNA polymerase subunits beta and beta’ (RpoB/RpoB’), and DNA polymerase I (PolA), representing fundamental cellular functions, and examined their distribution in the clusters. We found these proteins to be remarkably conserved. Although the *groEL* gene was universally conserved in all the organisms in the study, the protein was not represented in all the deduced proteomes. The genes for RpoB and RpoB’ were missing from two genomes and merged in 88 genomes, and the sequences were sufficiently divergent that they formed separate clusters for 18 RpoB proteins (seven clusters) and 14 RpoB’ proteins (three clusters). For PolA, 52 organisms lacked an identifiable sequence, and seven sequences were sufficiently divergent that they formed five separate clusters. Interestingly, organisms lacking an identifiable PolA and those with divergent RpoB/RpoB’ were almost all endosymbionts. Furthermore, we present a range of examples of annotation issues that caused the deduced proteins to be incorrectly represented in the proteome. These annotation issues represent a significant obstacle for high throughput analyses.

## Introduction

The concept of a minimal genome suggests the existence of a minimal set of genes for an organism to independently perform cellular functions (Koonin 2000; Koonin 2003; Gil et al. 2004). In search of a minimal gene set, researchers have artificially created organisms such as JVCI-syn1.0 and JVCI-syn3.0 that can grow in the laboratory with as few as 473 genes (Hutchison et al. 2016). The wealth of genome data currently available from bacterial organisms and the development of powerful clustering technology has enabled us to analyze millions of protein sequences from thousands of deduced proteomes in a single computational process. By comparing deduced proteomes from a diverse set of microorganisms we can gain insight into the distribution of putative minimal genes/proteins and ask beguilingly simple questions about protein conservation – for example, is a set of proteins conserved and if not, why not? For surely, in light of the minimal genome concept, all bacteria must contain some set of proteins essential for carrying out basic cellular functions. Clearly, the alternative hypothesis would be that a function is retained, but the sequence of the protein has diverged sufficiently that it no longer clusters into just one similarity group or that the function is performed by redundant proteins that are non-similar.

In this study, we examine the distribution and the level of conservation among four genes/proteins in Proteobacteria. To accomplish this, we used novel clustering technology to create clusters from the deduced proteomes of 2,307 complete Proteobacterial genomes encompassing 8.76 M protein sequences. Using the JVCI-syn3.0 minimal genome (Hutchison et al. 2016) to guide selection of proteins for analysis, we selected the chaperonin GroEL from the protein processing, folding, and secretion category (GroEL); DNA-directed RNA polymerase subunit beta and beta’ from the RNA metabolism category (RpoB/RpoB’); and DNA polymerase I from the DNA replication category (PolA). While these proteins are prevalent in Proteobacteria, their precise distribution and conservation are not known.

Our analysis revealed a remarkable degree of conservation in some proteins and different evolutionary paths in others. GroEL appeared to be the most conserved protein among the four – all Proteobacterial GroEL were clustered in just one cluster. PolA was also highly conserved. However, this was the only protein of the four that was missing from a large group of Proteobacteria which was almost completely comprised of endosymbionts. RpoB and RpoB’ exhibited behavior not observed in the other two – while highly conserved, these proteins created species-specific clusters suggesting protein differentiation.

Initially we were surprised that none of the clusters we examined contained representatives of all genomes in the study. However, on closer examination we learned that they should have, but due to two different types of annotation issues the deduced proteome was not complete. Our analysis highlighted two frequent problems in the annotation of complete genomes: protein misannotation and sequence discrepancies, the latter either real or technical. Of course, in both cases human error was also a factor. These annotation errors present a significant hindrance to investigators working with large datasets. Detecting errors in annotations is important because they have a tendency to propagate and, indeed, this propagation is increasing the number of errors with time (Schnoes et al. 2009). We suggest that our method of clustering can be used to substantially improve the quality of protein and genome annotation.
Results

## Results

### Microorganism data

At the time of this study, the total number of available Proteobacterial genomes in various stages of completion was 29,652. However, only 2,358 were marked as “complete.” Furthermore, 32 of the complete genomes were not accessible for download and an additional 19 genomes were disqualified from analysis for various reasons (Supplemental Table S1). The final set included 2,307 genomes comprising almost 8.76 M protein sequences (Supplemental Table S2).

In the final set, *γ*-Proteobacterial species accounted for nearly half of all complete genomes with the rest of the bacteria almost evenly split among α-, β-, and Δ/ε-Proteobacteria (Table 1). Members of the **Enterobacteriaceae** family comprised almost a quarter of the Proteobacteria. This fact is not surprising because this family contains many important human and animal pathogens and, as a result, it has been more intensively studied. Most of the protein sequences were located on chromosomes, but a significant number (269,461) were found on plasmids.

**Table 1.** Distribution of Major Proteobacterial Classes in the Study and Number of Protein Sequences.

| *Proteobacterial class* | *# of organisms* | *# of proteins* |
| --- | --- | --- |
| α-Proteobacteria | 380 | 1,272,409 |
| β-Proteobacteria | 332 | 1,514,315 |
| $\Delta/\epsilon$-Proteobacteria | 297 | 693,730 |
| γ-Proteobacteria | 1,298 | 5,210,577 |
| *Total* | 2,307 | 8,691,031 |

The genome sizes ranged from 0.11 Mb for *“Candidatus* (*Ca.*) Nasuia deltocephalinicola” to 14.78 Mb for *Sorangium cellulosum* str. So0157-2. The distribution of lengths was not uniform as shown in Fig. 1 which shows two prominent peaks corresponding to the most intensively studied pathogens - *Campylobacter* and *Helicobacter* spp. (peak at ~1.7 Mb) and *Salmonella* and *E. coli* spp. (peak at ~4.8 Mb).

**Figure 1:**
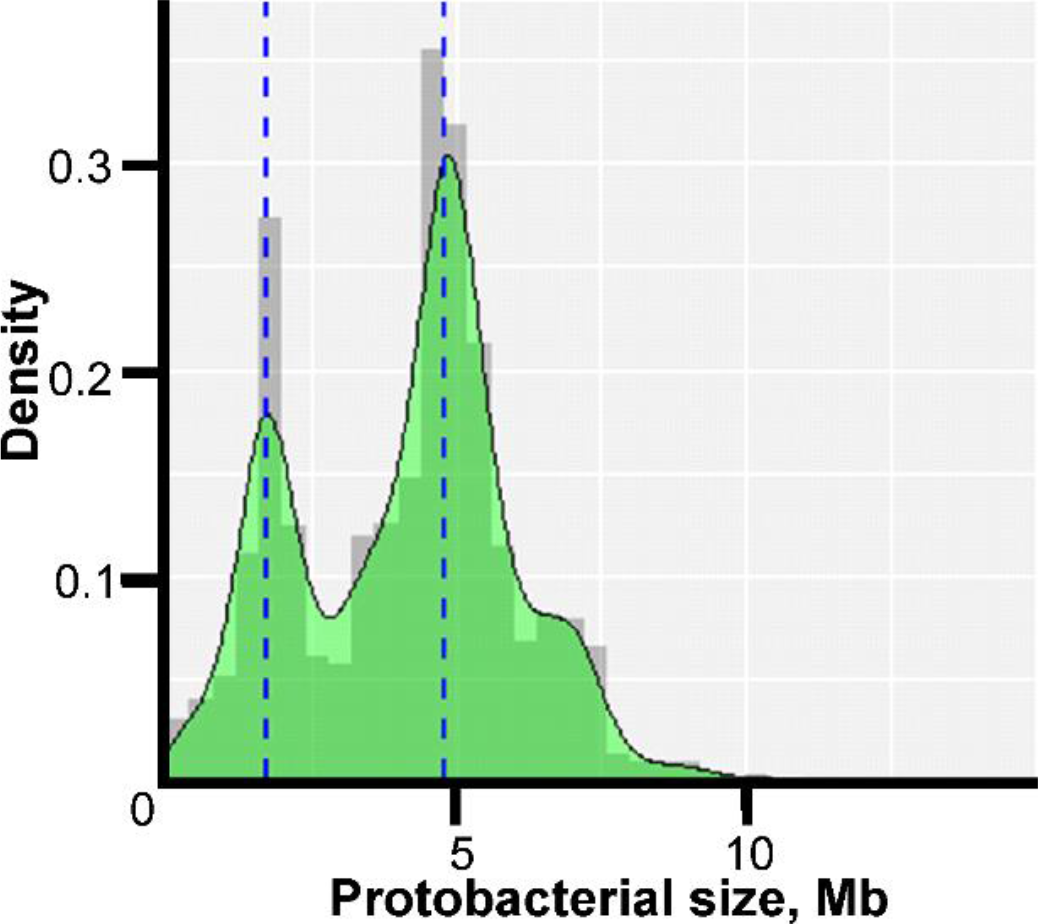
Density Histogram of Microorganism Lengths. The x-axis shows the genome size in Mb, and the y-axis shows the probability density function estimated from the histogram. The bimodal peaks occur because of the large number of Enterobacteraceae genomes (~4.8 Mb) and Campylobacteria and Helicobacteria genomes (~1.7 Mb) in the study.

The GC content varied from 13.5% for “*Ca.* Zinderia insecticola” str. CARI to almost 75% for the members of *Anaeromyxobacter* spp. (Supplemental Table S2). All major Proteobacterial classes appeared to be spread approximately evenly across the full range of GC content (Fig. 2). Higher GC content is associated with higher variability in genome size, perhaps allowing broader niche adaptation, while the genome size of Proteobacteria with low GC content varies much less.

**Figure 2:**
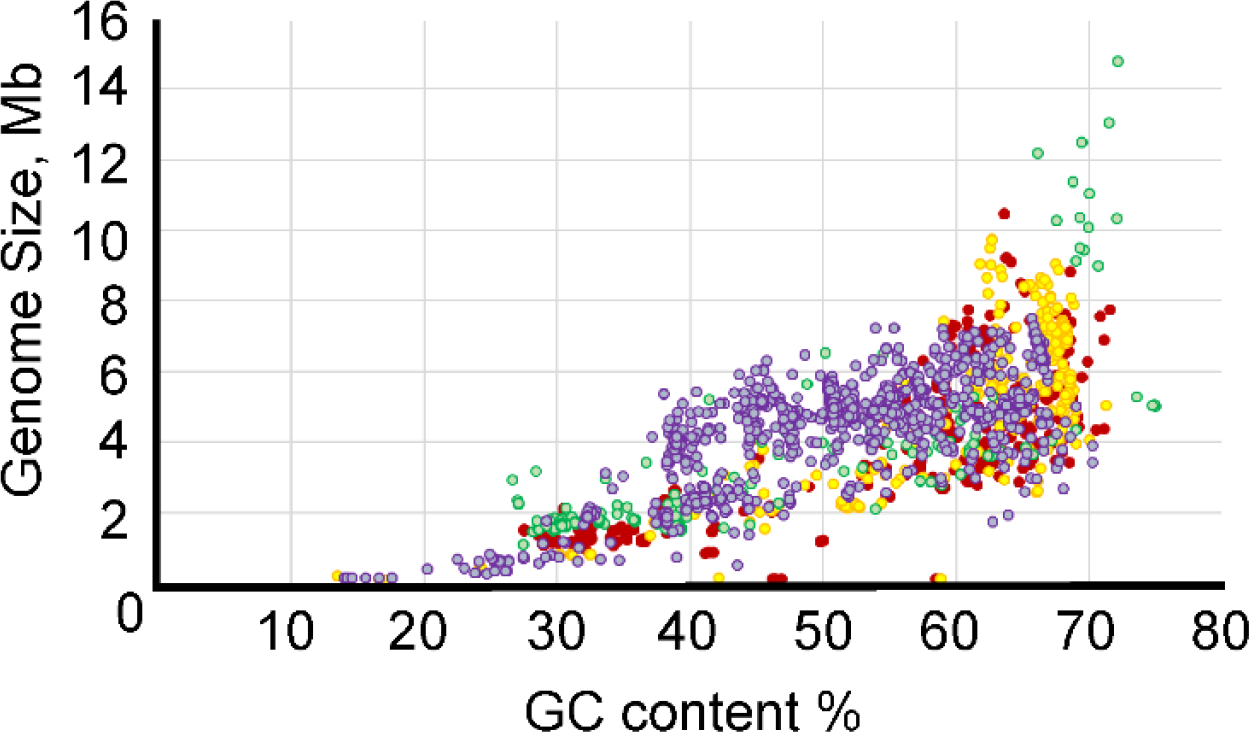
Relationship between GC Content and Genome Length. The x-axis shows GC content, and the y-axis shows genome size. The color codes are as follows: red - α-Proteobacteria; yellow - β-Proteobacteria; purple - γ-Proteobacteria; and green - δ/ε-Proteobacteria.

Clustering resulted in 707,311 singleton and non-singleton clusters. A singleton is a protein sequence that does not align with any other sequence in the set. These sequences are unique not only among the organisms, but also within an organism. There were 472,820 such sequences or, equivalently, 5.4% of all sequences were singletons. There were also 10,049 clusters in which there was more than one sequence, but the sequences in each cluster were from the same organism. The total number of nonsingleton clusters was 224,442.

The 224,442 clusters of millions of similar protein sequences, ranging in size from 2 to 2,894 sequences, represent a vast amount of cached information. As a first step in analyzing these clusters, we examined the representation of the organisms used in the study in each cluster. We found, for example, that no cluster contained a sequence from every organism in the study and that only 0.1% of the clusters contained sequences from at least 90% of the organisms. We decided to explore the presence of four putatively broadly conserved genes/proteins in the clusters. Our selection was guided by a recent paper published on a minimal genome (Hutchison et al. 2016), and our choice was based on a desire to have three different functional classes of proteins.

The core essential genes that support cellular life can be partitioned into six functional classes (Kogoma 1997; Smalley et al. 2003; Gil et al. 2004; Friedberg et al. 2005; Sydow and Cramer 2009): 1) DNA metabolism including DNA replication, repair, and modification; 2) RNA metabolism including translation and RNA degradation; 3) protein processing, folding, and secretion including posttranslational modification and translocation; 4) cellular processes including cell division and substrate transport; 5) energetic and intermediary metabolism including glycolysis and lipid metabolism; and 6) a small set of conserved but poorly characterized genes. The four proteins that we chose for our study (GroEL, RpoB/RpoB’, and PolA) correspond to the first three categories.

During our analysis we uncovered various protein and gene annotation issues that required manual inspection and presented significant hindrance to our study. Most frequently we encountered fragmented proteins and, in a few cases, a gene was simply overlooked during annotation. Frameshifts can result in deduced protein fragmentation. These frameshifts can be either real or due to sequencing error. With high-throughput sequencing technology, frameshifts can be introduced when the incorrect number of residues are recorded in polynucleotide tracks (Henson et al. 2012). Protein fragmentation can also occur when a stop codon is introduced into a coding sequence. Annotation problems caused apparent absence of proteins in the deduced proteomes, and fragmented proteins often clustered incorrectly because of their small size. Collectively, we refer to these as “misannotations” and discuss them individually in each section.

## GroEL

### Protein background

Chaperones are proteins that help in intermolecular assembly without being part of the final product (Fenton and Horwich 2003). Chaperonins are a class of molecular chaperones that assist other proteins to fold, and they act on fully synthesized proteins (Bhutani and Udgaonkar 2002; Chapman et al. 2006; Apetri and Horwich 2008). GroEL belongs to Group I of the chaperonin family (Fenton and Horwich 2003). It is characterized by a ring-shaped structure consisting of seven identical subunits (Fenton and Horwich 2003; Lin et al. 2008). The work by Lin *et al.* showed that GroEL stimulates protein folding through forced unfolding rather than acting as a passive aggregation inhibitor (Lin et al. 2008).

Our analysis shows that GroEL is one of the most conserved proteins among Proteobacteria. All GroEL sequences are found in just one cluster (Cl. 3128) indicating high conservation of sequence and biological function. To function correctly, GroEL requires its accessory protein, the co-chaperonin GroES (Lin et al. 2008). We found remarkably conserved synteny of the *groEL/groES* system. The two protein coding genes are always positioned next to each other with *groES* immediately upstream from *groEL*, and both are transcribed in the same direction.

### GroEL statistics and clusters

From the 2,307 genomes in the study, we detected 2,830 *groEL* coding sequences. At first glance not all organisms had a GroEL representative in the main GroEL cluster (Cl. 3128). Upon delving into the genome sequences, we ascertained that all the genomes in our study contained a *groEL* sequence. However, as described below, two were not annotated at all, and two contained frameshifts within the gene and were annotated as pseudogenes and, thus, these four were not included in the proteome.

The conservation of GroEL is evident from its presence in all the genomes in the study to the conservation of its length. Although there were some outliers, the standard deviation was small – only nine amino acids (aa) with 94% of GroEL proteins (2,646 out of 2,826) between 535 and 550 aa.

Only 3% of the *groEL* genes are on plasmids; the rest are located on chromosomes. The vast majority of microorganisms (2,254) maintain their *groEL* on the chromosome, 52 microorganisms have *groEL* sequences on a plasmid as well as on the chromosome while only one organism – *Methylobacterium aquaticum* – has its sole *groEL* on a plasmid. The multiplicity of *groEL* is higher for plasmids than for chromosomes – 1.66 versus 1.19 – indicating that on average there are more *groEL* genes per plasmid than per chromosome.

Of the 2,307 organisms studied, 1,948 [^1^ This number includes missannotated GroEL, but excludes partial/dysfunctional GroEL.] have only 1 *groEL* gene (Supplemental Fig. S1). However, there are a few Proteobacteria with a much higher multiplicity of *groEL.* Among them, *Bradyrhizobium diazoefficiens* str. USDA 110 and *Sinorhizobium meliloti* str. AK83 and SM11 stand out with the most *groEL* genes, seven. Members of the genus *Bradyrhizobium* have at least three *groEL* genes with an average of 4.9 genes per strain. Similarly, genus *Sinorhizobium* has four or more *groEL* with an average of 5.7. All members of this genus allocate *groEL* sequences approximately evenly between plasmids and chromosomes with one exception – *S. meliloti* str. AK83 contains all its *groEL* genes on the chromosome.

### GroEL annotation issues

There are a few outliers in length among GroEL proteins – 42 sequences were found that were much smaller than the typical GroEL. There were two reasons for these smaller sequences: 1) when multiple copies of the gene are present, occasionally one copy might become truncated, or pseudogenized, and 2) if the sequence contained a frameshift, the annotators handled this in different ways – sometimes they annotated the larger part of the gene as a CDS, sometimes they annotated both parts, and sometimes they just annotated the gene but not its product, or missed the gene entirely, in which case there was no representation in the proteome. An example of case 1: *Pseudomonas putida* (Org. 1316) has two GroEL sequences one of which is significantly shorter than the other – 292 aa vs. 546 aa (CDS AHZ77123.1 and AHZ79220.1). However, the shorter GroEL is most likely dysfunctional because it is approximately half the length of a typical GroEL protein. In terms of organismal function, this doesn’t create a problem because the full length GroEL is still available.

In an example of both cases 1 and 2, a similar yet different situation occurred with *Bradyrhizobium diazoefficiens* strain NK6 (Org. 2290). This organism has five intact *groEL* genes; the sixth gene appears to have become dysfunctional with both an internal stop codon and a frameshift (Fig. 3a). All the gene segments were annotated as GroEL; however, only the largest and most 3′ part of the gene (encoding CDS BAR62159.1) falls in the main GroEL cluster (Cl. 3128). As an example of case 2, “*Ca.* Midichloria mitochondrii” str. IricVA (Org. 713) was annotated as having only one *groEL* gene encoding a product of 385 aa (protein AEI89468.1). Upon closer inspection of the genomic region, coding for *groEL* revealed that there is a frameshift, and the remaining portion of the gene is present but not annotated (Fig. 3b). We cannot know if this frameshift is a sequencing error or is a real frameshift creating a shortened protein, but this is an important distinction as this is the only GroEL in this organism. In this particular case, this shortened version of GroEL may be long enough to be functional. In a unique case of a “short” GroEL, upon examination of the genome sequence of *Burkholderia glumae* str. PG1 (Org. 1674), the annotated CDS (AJK45248.1) is only 481 aa, but the genomic region 5′ to the annotated start codon is open and corresponds to the 5’ portion of the *groEL* gene (Fig. 3c). This gene is, in fact, misannotated, and if the start codon at position 816988 is used, a full length GroEL of 546 aa is present in this locus. In this latter case, a short sequence is included in the protein annotation; however, this is an annotation error.

**Figure 3:**
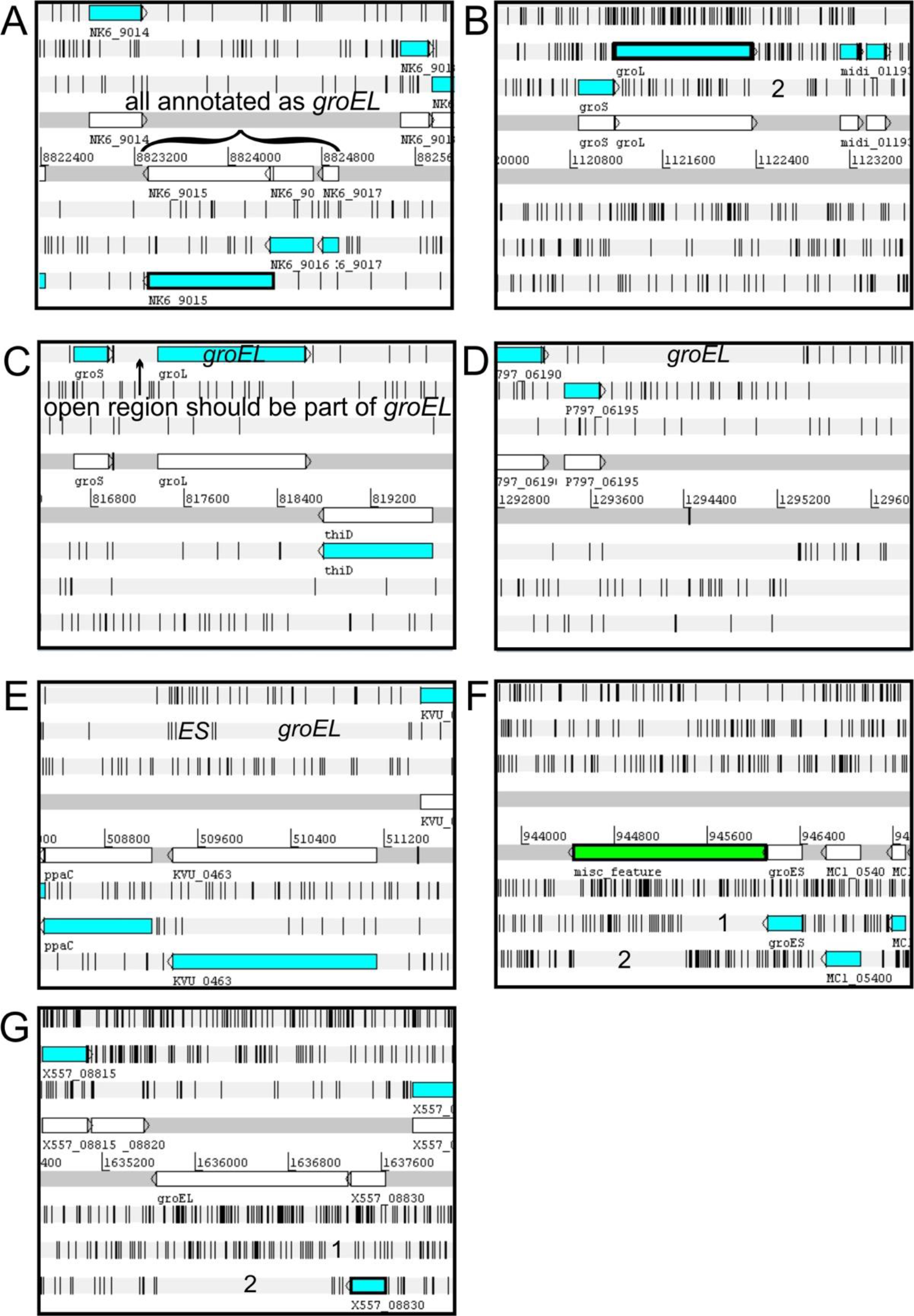
Annotation Issues Detected for *groEL.* Images captured from Artemis genome browser. The three forward reading frames are shown on top and three reverse reading frames on bottom as horizontal light gray bars. Black vertical lines on the light gray bars indicate stop codons in the six reading frames. The two horizontal dark gray bars contain gene annotations in white. The blue bars are the annotated CDSs, or protein coding genes. Numbers along the middle of the image indicate base position in the genome, while letters and numbers near a gene indicate the locus identifier. A) *Bradyrhizobium diazoefficiens* strain NK6 contains a *groEL* gene with a stop codon and a frameshift. All segments were annotated as GroEL and were incorporated into the proteome. B) In “*Ca*. Midichloria mitochondrii” str. IricVA the *groEL* gene contains a frameshift that was not recognized, and the first part of the gene was annotated as GroEL and was incorporated into the proteome as a truncated protein. C) For *Burkholderia glumae* str. PG1, a truncated *groEL* gene and thus protein were annotated although the region upstream from the utilized start codon corresponds to *groEL* and has a start codon in the appropriate position to encode a full length protein. D) *Pseudomonas aeruginosa* str. VRFPA04 was simply missing annotation for the *groEL* gene. E) As a potential example of algorithmic error, in *Ketogulonicigenium vulgare* str. WSH-001, the gene for *groEL* was overlooked in favor of a hypothetical gene on the opposite strand. F) In *Rickettsia parkeri* str. Portsmouth the *groEL* gene was recognized as containing a frameshift and was annotated as a miscellaneous feature. G) In *Francisella tularensis* subsp. holarctica str. PHIT-FT049, the frameshift in *groEL* was also recognized; however, the annotation was different than in F) as it was annotated as a pseudogene. In cases A)-C) aberrant deduced protein species were incorporated into the deduced proteome, while in cases D)-G) no protein products were incorporated into the deduced proteome.

Other annotation errors were found leading to no representation of GroEL in the deduced proteome. Two genomes were found to contain a full-length, intact *groEL* sequence even though none had been annotated: 1) *Pseudomonas aeruginosa* str. VRFPA04 (Org. 1101) (Fig. 3d) and 2) *Ketogulonicigenium vulgare* str. WSH-001 (Org. 730) (Fig. 3e). In the latter case, a hypothetical gene was annotated on the opposite strand. In two other genomes the *groEL* gene contained a frameshift and was annotated differently in each. For example, *Rickettsia parkeri* str. Portsmouth (Org. 878) region complement (944449…946091) is marked as “misc_feature” (Fig. 3f), while *Francisella tularensis* subsp. holarctica str. PHIT-FT049 (Org. 1197) has the *groEL* gene annotated with the qualifier “pseudo” and comment “disrupted” (Fig. 3g).

From the analysis above, we conclude that all of the organisms in our study contained at least one gene encoding the GroEL protein. However, not all of these genes were translated into the deduced proteome because of annotation errors or frameshifts that may be real or due to sequencing errors.

## DNA-dependent RNA polymerase subunits β and β’

### Protein background

DNA-dependent RNA polymerases, found in all domains of life, are essential for cellular function. RNA polymerases catalyze the transcription of DNA into RNA and facilitate nucleotide attachment and RNA elongation (Berg et al. 2002). They also normally have proofreading and termination recognition capability (Sydow and Cramer 2009). A very large molecule is required to perform such a wide variety of functions. Indeed, a complete RNA polymerase holoenzyme consists of six subunits (Helmann and Chamberlin 1988) of which RNA polymerase subunits β’ (RpoB’) and β (RpoB) are the largest and second largest. These are encoded by the *rpoC* and *rpoB* genes, respectively (Helmann and Chamberlin 1988).

Initially only RpoB was considered for analysis. However, it soon became clear that RpoB’ must also be included for two reasons: often the annotations were switched, and while the majority of microorganisms have separate RpoB and RpoB’, *Helicobacter* and *Wolbachia* spp. encode these two proteins as a single polypeptide. Just as in the case of the *groEL/groES* system, we observed highly conserved synteny with the *rpoB/rpoC* system. The two protein coding genes are positioned next to each other with *rpoC* immediately upstream from *rpoB.* Both are usually transcribed in the same frame with very few instances of the genes occurring in different frames.

### Protein statistics and clusters

Most Proteobacterial RpoB and RpoB’ were found in two main clusters – Cls. 3025 and 3026, respectively. The mean length of RpoB was 1,360 aa while that of RpoB’ was 1,412 aa (Tables 2 and 3). RpoB’ is usually considered to be longer than RpoB, and we found 2,060 sequences of RpoB’ that were longer than RpoB with the largest difference being 126 aa. However, the situation was reversed for 137 microorganisms for which RpoB’ was shorter than RpoB with the largest difference being 232 aa.

**Table 2.** RpoB Cluster Details.

| Cluster / Organism<sup>a</sup> | # sequences | Min | Mean | Max | Std dev | Notes |
| --- | --- | --- | --- | --- | --- | --- |
| 3025 | 2188 | 1063 | 1360.8 | 1499 | 20.6 | main RpoB cluster |
| 49264<sup>b</sup> | 8 | 1258 | 1262.8 | 1267 | 3.7 | "*Ca. Carsonella ruddii*"<sup>b</sup> |
| 201571<sup>b</sup> | 2 | 1342 | 1342.5 | 1343 | 0.7 | "*Ca. Nasuia deltocephalinicola*"<sup>b</sup> |
| 211635<sup>b</sup> | 3 | 1205 | 1223 | 1235 | 15.9 | "*Ca. Hodgkinia cicadicola*"<sup>b</sup> |
| 174727<sup>b</sup> | 2 | 1290 | 1290 | 1290 | 0 | "*Ca. Tremblaya princeps*"<sup>b</sup> |
| Singleton Org. 357<sup>b</sup> | 1 | -- | 1267 | -- | -- | "*Ca. Hodgkinia cicadicola*" Dsem<sup>b</sup> |
| Singleton Org. 104<sup>b</sup> | 1 | -- | 1291 | -- | -- | "*Ca. Tremblaya phenacola*" PAVE<sup>b</sup> |
| Singleton Org. 519<sup>b</sup> | 1 | -- | 1291 | -- | -- | "*Ca. Zinderia insecticola*" CARI<sup>b</sup> |

<sup>a</sup>Microorganism name is given with its number if it's RpoB is a singleton.
<sup>b</sup>indicates endosymbionts.

**Table 3.**
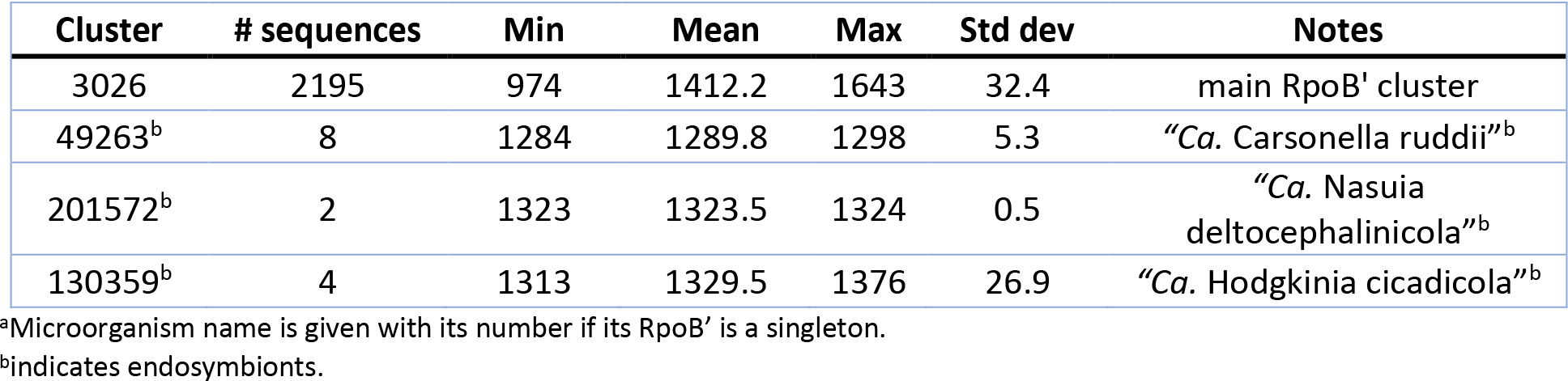
RpoB’ Cluster Details.

| Cluster | # sequences | Min | Mean | Max | Std dev | Notes |
| --- | --- | --- | --- | --- | --- | --- |
| 3026 | 2195 | 974 | 1412.2 | 1643 | 32.4 | main RpoB' cluster |
| 49263<sup>b</sup> | 8 | 1284 | 1289.8 | 1298 | 5.3 | "*Ca. Carsonella ruddii*"<sup>b</sup> |
| 201572<sup>b</sup> | 2 | 1323 | 1323.5 | 1324 | 0.5 | "*Ca. Nasuia deltocephalinicola*"<sup>b</sup> |
| 130359<sup>b</sup> | 4 | 1313 | 1329.5 | 1376 | 26.9 | "*Ca. Hodgkinia cicadicola*"<sup>b</sup> |

<sup>a</sup>Microorganism name is given with its number if its RpoB' is a singleton.
<sup>b</sup>indicates endosymbionts.

Among 2,307 Proteobacteria, 2,217 (96%) encoded their RpoB and RpoB’ separately, and 88 microorganisms encoded merged RpoB/RpoB’ proteins (Table 4), but two of the 88 organisms contained a frameshift in the merged *rpoB/rpoC* gene (Orgs. 941 and 1222). Two strains of “*Ca.* Filomicrobium marinum” did not have identifiable RpoB or RpoB’ (Supplemental Table S3). Multiplicity is a very rare event for RpoB and RpoB’. Three microorganisms duplicated their *rpoB* and *rpoC* genes (Orgs. 218, 510, and 1981), and one microorganism, *Thioploca ingrica*, had two copies of *rpoC* but only one *rpoB.*

**Table 4.** Merged RpoB/RpoB’ Cluster Details.

| Cluster | # sequences | Min | Mean | Max | Std dev | Notes |
| --- | --- | --- | --- | --- | --- | --- |
| 26658 | 79+1<sup>a</sup> | 2296 | 2879.9 | 2894 | 67.8 | *Helicobacter* spp. |
| 26658 | 8 | 2837 | 2838.5 | 2842 | 1.7 | *Wolbachia* spp.<sup>b</sup> |

<sup>a</sup>+1 indicates Org. 1222. Although its protein annotation is different, its gene indicates it should belong to this cluster (see text).
<sup>b</sup>indicates endosymbionts.

Only four Proteobacterial strains carried their *rpoB* and *rpoC* genes on plasmids and did so in pairs., i.e., if *rpoB* was on a plasmid then *rpoC* was there as well. All plasmid *rpoB* and *rpoC* were exceptions in their respective species. For example, only one strain of *Shewanella baltica* OS155 placed its *rpoB/rpoC* system on a plasmid; the remaining six *S. baltica* strains had it on the chromosome. The same situation occurred in *Serratia liquefaciens* str. ATCC 27592 and *Salmonella enterica* subsp. enterica serovar Senftenberg. *Pseudomonas chlororaphis* str. PCL1606 (Org. 1775) had its *rpoB* on a plasmid; however, a hypothetical gene was annotated on the opposite strand instead of its *rpoC* as in Fig. 3e. The rest of the microorganisms contained their *rpoB* and *rpoC* on chromosomes.

### RpoB and RpoB’ clusters from Endosymbionts

RpoB and RpoB’ from some endosymbionts formed their own clusters. For example, they grouped into Cls. 49264 and 49263 for all members of “*Ca.* Carsonella ruddii” (Tables 2 & 3). The same circumstance occurred for endosymbionts “*Ca.* Nasuia deltocephalinicola” (RpoB – Cl. 201571, RpoB’ – Cl. 201572) and “*Ca.* Hodgkinia cicadicola” (RpoB – Cl. 211635 and singleton, RpoB’ – Cl. 130359). “*Ca.* Tremblaya” spp. had their RpoB sequences in a separate cluster and a singleton, but their RpoB’ proteins were in the main RpoB’ Cl. 3026 (Tables 2 & 3).

RpoB and RpoB’ from endosymbionts were usually shorter than those in Cls. 3025 and 3026, but they preserved the domains necessary to perform the required functions. However, because of low sequence identity (<40%) with RpoB/RpoB’ sequences in the main clusters, separate clusters were formed (Tables 2 and 3).

### RpoB and RpoB’ of *Helicobacter* and *Wolbachia* spp

Members of *Helicobacter* and *Wolbachia* spp. had a separate cluster for RpoB/RpoB’ (Table 4) because they encoded a merged protein with an average length of 2,876 aa. This is a unique situation among Proteobacteria. There were 79 complete RpoB/RpoB’ sequences from *Helicobacter* spp. and eight sequences from *Wolbachia* spp. (Table 4). *Helicobacter* spp. had only one strain, *H. pylori* str. SouthAfrica20 (Org. 1222), that appeared to have separate *rpoB* and *rpoC* genes; however, the *rpoB* gene contained a frameshift (the addition of a single A residue in the polynucleotide tract upstream from the stop codon in *rpoB* would result in a single gene encoding the multifunctional *rpoB/rpoC*). *H. pylori* str. Rif1 (Org. 941) also contained a frameshift in the merged gene for RpoB/RpoB’; however, the larger part of the protein was 2,296 aa and fell into Cl. 26658 while the amino terminal part of 594 aa was in the main RpoB cluster (Cl. 3025). If there were no frameshift, these protein segments would have created a 2,890 aa protein consistent with the lengths of the other *H. pylori* merged RpoB/B’ proteins.

### RpoB and RpoB’ annotation issues

Most of the annotation issues described for GroEL were encountered with RpoB and RpoB’, which resulted in partial or complete omission of these gene products from proteomes. In total, 25 Proteobacterial genomes had annotation issues including five with omitted RpoB’, four with omitted RpoB, one with omitted RpoB and only partial RpoB’ (*Salmonella enterica* subsp. enterica serovar Enteritidis str. EC20120916, Org. 1265), ten with only parts of RpoB or RpoB’ annotated, and another five with both RpoB and RpoB’ entirely omitted (Supplemental Table S3). As before, most partial or complete omissions were results of frameshifts. We noted that in every case where the *rpoB* or *rpoC* product was omitted, it was an exception in the respective species. For example, every genome among 75 *Vibrio* strains had RpoB and RpoB,’ yet *Vibrio cholerae* str. LMA3984-4 (Org. 637) had neither. A closer inspection showed a genomic region with annotated genes containing several frameshifts, although one was annotated incorrectly (locus tags VCLMA_A0287, VCLMA_A0288). The sequencing of this genome was done using short DNA reads – a highly error prone method (Chaparro et al. 2011). In fact, the strain had 196 frameshifts. A similar situation occurred for *Advenella kashmirensis* str. WT001 (Org. 718) whose genome had 925 frameshifts.

Of 2,307 microorganisms, only strains of “*Ca.* F. marinum” (Orgs. 1822 and 1823) did not have identifiable RpoB and RpoB’. These were the only two strains of this species with fully sequenced genomes. Their closest related species is *F. insigne* (NCBI accession GCA_900104305.1) whose genome is not fully sequenced. Alignment of the genomic fragment containing *rpoB* and *rpoC* against the “*Ca.* F. marinum” genomes revealed an absence of synteny and poor sequence similarity. Similar results were obtained by aligning “*Ca.* F. marinum” genomes against a related *Rhodomicrobium* genome (results not shown). The question remains open whether these “*Ca.* F. marinum” strains have RpoB or RpoB’. From the analysis above, it is clear that RpoB and RpoB’ are well conserved but more diverse than GroEL. Only two genomes were missing genes for these proteins, and both were for the same species. A small subset of genomes was found to have a merger of the two genes *rpoB* and *rpoC* that encoded a bifunctional protein. For both RpoB and RpoB’ there were some sequences that varied sufficiently to be grouped in different clusters. Interestingly, all the variable protein sequences that grouped separately from the two main clusters belonged to endosymbionts and were slightly shorter in length.

## DNA Polymerase I

### Protein background

DNA polymerases are a diverse family of enzymes that participate in DNA replication and repair (Cooper 2000). In prokaryotes, five types of DNA polymerase have been identified – Pol I-V. Pol I, which we will refer to by its more commonly used name PolA, is one of the most abundant polymerases, e.g., in *E. coli* it is responsible for >95% of polymerase activity (Camps and Loeb 2004). The primary function of PolA is DNA repair during the replication process (Sutton and Walker 2001; Friedberg et al. 2005; Hübscher et al. 2010), and the structure of a typical prokaryotic PolA supports this functionality (Bernad et al. 1989).

A canonical PolA consists of two linked functional subunits – PolA.1 and PolA.2 – containing the following three domains listed in order from amino-terminus to carboxy-terminus (Bernad et al. 1989): 5’ to 3’ exonuclease (PolA.1), 3’ to 5’ exonuclease, and 5’ to 3’ polymerase (PolA.2) (Fig. 4). The majority of microorganisms in our data set (2,231 or ~97%) contained a complete PolA with both subunits PolA.1 and PolA.2. In the rare cases when a frameshift was present, this usually resulted in separation of the PolA.1 and PolA.2 domains. In this study we considered any PolA without one of the three domains to be an incomplete PolA.

**Figure 4:**
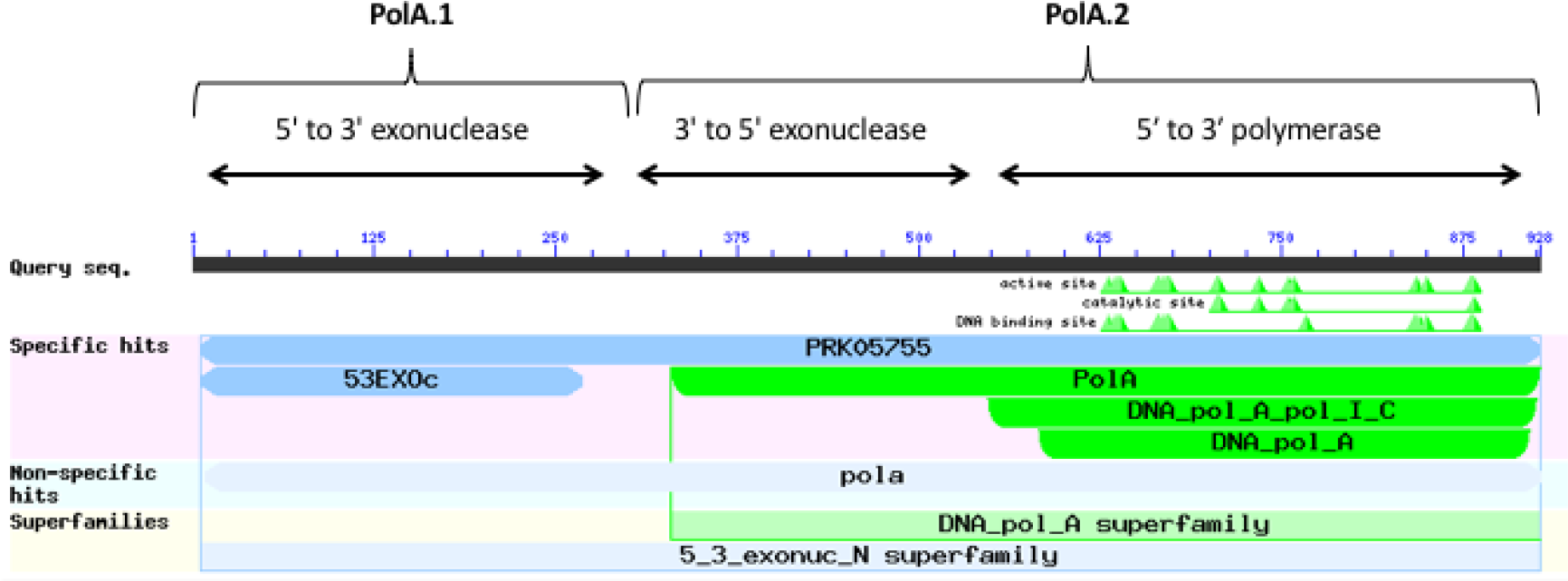
Domain Architecture of PolA. Image captured from NCBI conserved domains showing results for a typical PolA. There are three domains in PolA: 5’ to 3’ exonuclease, 3’ to 5’ exonuclease, and 5’ to 3’ polymerase. The 5’ to 3’ exonuclease domain (PolA.1) is often separated from the other two domains (PolA.2).

### Protein statistics and clusters

Together there were 2,258 complete PolA sequences. The average PolA was slightly longer than 900 aa (Table 5). For proteins longer than 950 aa, there was extra sequence inserted either before or after the PolA.1 domain. Five microorganisms – members of *Azospirillum spp., Methylobacterium spp.*, and *Salmonella enterica* subsp. enterica serovar Senftenberg – allocated their only *polA* gene on plasmids, one bacterium *Methylobacterium extorquens* str. AM1 had *polA* on its plasmid and chromosome, while the rest of the microorganisms had their *polA* genes on chromosomes. Multiplicity was a rare event for *polA.* Only four microorganisms had duplicate *polA* genes, three of which belonged to the delta/epsilon class.

**Table 5.** PolA Cluster Details.

| Cluster / Organism<sup>a</sup> | # sequences | Min | Mean | Max | Std dev | Notes |
| --- | --- | --- | --- | --- | --- | --- |
| 2948 | 2241 | 588 | 922.5 | 1165 | 35.75 | main cluster |
| 168070 | 2 | 759 | 780 | 801 | 29.70 |  |
| 210955 | 2 | 264 | 288 | 312 | 33.94 | partial PolA annotated |
| Singleton<br>Org. 1970 | 1 | -- | 863 | -- | -- | *Labilithrix luteola* |
| Singleton<br>Org. 1172 | 1 | -- | 912 | -- | -- | "*Ca. Babela massiliensis*" |
| Singleton<br>Org. 475 | 1 | -- | 537 | -- | -- | *Sorangium cellulosum* So<br>ce56 |
| Multiple spp. | 10 | -- | -- | -- | -- | No PolA annotated<sup>b</sup> |

<sup>a</sup>Microorganism name is given with its number if its PolA is a singleton.
<sup>b</sup>Detailed information in Table 6.

There were two clusters (Cls. 2948 and 168070) and two singletons (Orgs. 1970 and 1172) of complete PolA protein sequences (Table 5). The majority of PolA sequences (2241) were in Cl. 2948 which predominantly contained complete PolA protein sequences (PolA.1+PolA.2) but also contained 11 incomplete sequences corresponding to PolA.2, ranging in length from 588 to 775 aa. However, these incomplete sequences had matching PolA.1 parts in other clusters because they had become separated due to a frameshift in the gene or because they had been missed during annotation.

The only organism for which we could not find an identifiable DNA polymerase I was *Mannheimia haemolytica* str. USMARC_2286 (Org. 1073). All other nine members of this species had a complete PolA with a consistent length of 952 aa. However, no identity was found when aligning the *Mannheimia haemolytica* str. USMARC_2286 genome against known *M. haemolytica* PolA using the tblastn module of BLAST (Altschul et al. 1990).

Cluster 168070 contained only two species – *Desulfurella acetivorans* str. A63 (Org. 1176) and *Hippea maritima* str. DSM 10411 (Org. 634). The two PolA sequences in this cluster were shorter than a canonical PolA due to a complete or partial loss of the 3’ to 5’ exonuclease domain (part of PolA.2) which is responsible for proofreading activity. Whether the loss of this domain impairs the normal function of PolA or other enzymes perform proofreading during DNA replication remains an open question.

*Labilithrix luteola* (Org. 1970) had two different types of PolA – a canonical PolA (945 aa) located in the main cluster (Cl. 2948) and a singleton PolA of 863 aa (Table 5). The latter not only lacked an identifiable 5’-3’ exonuclease (PolA.1) but also a 3’-5’ exonuclease. Whether this strain of *L. luteola* has new, previously unknown types of 3’-5’ and 5’-3’ exonucleases or this is just a dysfunctional PolA can be determined only experimentally.

The singleton PolA of “*Ca.* Babela massiliensis” (Org. 1172) was similar to the shorter PolA of *L. luteola* in that it lacked an identifiable 5’-3’ exonuclease, but its amino acid composition was very different not only from PolA of *L. luteola* but also from the PolA’s in any other cluster. “*Ca.* B. massiliensis” is a representative of an amoeba parasite (Pagnier et al. 2015). Low sequence similarity can explain why this PolA is a singleton; typical sequence identity calculated by BLAST when aligning with members of other clusters was 20%-30%.

### Endosymbionts

PolA was the only protein in our study that was absent from a large group of microorganisms, which were almost all endosymbionts. Among 81 endosymbionts representing 26 species, 29 endosymbionts had complete PolA, 23 had only PolA.1, and the remaining 29 had no identifiable PolA or part of PolA (Supplemental Table S4). We note that *Buchnera aphidicola* APS was one of the first fully sequenced genomes (Shigenobu et al. 2000; Tamas et al. 2002; van Ham et al. 2003), and the incomplete PolA, the 5’-3’ exonuclease (PolA.1), was annotated incorrectly as DNA polymerase I (Gil et al. 2004). As a result, successive strains of *B. aphidicola* were annotated in the same manner even though their average length is only 290 aa. Our analysis showed that among 18 members of *B. aphidicola*, only Orgs. 485 and 702 have full length PolA in Cl. 2948, the major PolA cluster. The remaining 16 *B. aphidicola* have only PolA.1. The situation differs for *Wolbachia spp.* for which all eight members have the full PolA protein.

All together we identified 81 organisms in our study as endosymbionts. Of these, almost 2/3 (52) lacked full-length DNA polymerase I (Supplemental Table S4). On the one hand, this could be expected because endosymbionts owe a significant part of their functional machinery to their hosts. On the other hand, it is surprising that PolA is missing, not a longer protein such as RpoB or RpoB’.

### Other DNA polymerase A family proteins

In addition to PolA, we found a number of other clusters with proteins annotated as DNA polymerase I. However, upon inspection these were found to be a separate set of DNA polymerase A family proteins.
A total of 143 sequences represented 108 species, and they were in addition to the canonical PolA protein (Supplemental Table S5). They were significantly shorter than the canonical PolA with an average length of only 626 aa. Their separate clustering indicates, perhaps, a different evolutionary path – some were annotated as phage-origin or phage-related DNA polymerase, but most were designated as conserved hypothetical proteins.

### PolA annotation issues

There were 23 microorganisms whose PolA contained either a frameshift, was partially annotated, or was entirely omitted from the proteome (Table 6). This includes eight Proteobacteria with a frameshift in PolA as in Fig. 3a, three organisms with a partial PolA annotated due to the incorrect choice of the start codon as in Fig. 3c, and ten bacteria with PolA absent from the proteome for various reasons (Supplementary Table S6). In the case of *Bordetella pertussis* (Org. 2300), *polA* does not have an associated protein file because although it is annotated, it contains seven stop codons.

**Table 6.** Misannotations and Frameshifts in *polA* Genes and Deduced Proteins.

| Organism Number | Organism Name | Misannotation Type<sup>a</sup> | Gene Annotation |
| --- | --- | --- | --- |
| 21 | *Escherichia coli* CFT073 | L | Absent |
| 390 | *Yersinia pestis* D182038 | J | Multiple stop codons |
| 550 | *Ketogulonicigenium vulgare* Y25 | A | Frameshift |
| 590 | "*Ca. Liberibacter solanacearum*" CLso-ZC1 | A | Frameshift |
| 660 | *Haemophilus influenzae* F3047 | C | Missed start codon |
| 975 | *Enterobacter aerogenes* EA1509E | A | Frameshift |
| 1085 | *Klebsiella pneumoniae* JM45 | D | Absent |
| 1105 | "*Ca. Pantoea carbekii*" | A | Frameshift |
| 1140 | *Hyphomicrobium nitrativorans* NL23 | G | Absent |
| 1148 | *Ehrlichia muris* AS145 | H | Gene annotated, protein not |
| 1212 | *Yersinia similis* | D | Absent |
| 1235 | *Escherichia coli* ST2747 | G | Absent |
| 1291 | *Azospirillum brasilense* | G | Absent |
| 1292 | *Dyella jiangningensis* | G | Absent |
| 1323 | *Halomonas campaniensis* | C | Missed start codon |
| 1474 | *Burkholderia thailandensis* E254 | B | Frameshift |
| 1587 | "*Ca. Ishikawaella capsulata*" Mpkobe | A | Frameshift |
| 1796 | "*Ca. Pantoea carbekii*" | A | Frameshift |
| 1819 | *Francisella tularensis* subsp. *tularensis* str. SCHU S4 substr. NR-28534 | A | Frameshift |
| 2177 | *Campylobacter jejuni* | G | Absent |
| 2278 | *Aggregatibacter actinomycetemcomitans* NUM4039 | A | Frameshift |
| 2288 | *Pseudomonas aeruginosa* | C | Missed start codon |
| 2300 | *Bordetella pertussis* | K | Absent |

<sup>a</sup>Refer to Supplemental Table S6 for annotation codes and Fig. 3 for examples.

## Protein misannotations

For each of the four proteins presented above, we highlighted issues involving annotation of their genes which resulted in truncated or missing proteins. In this section, we focus on just one of the four to highlight issues with gene product annotation, i.e., protein nomenclature. Both can be considered as types of misannotation, but they are distinctly different. The former often arises from sequence discrepancies – either real or technical – but can also be due to algorithmic or human error. The latter is often due to confusion, lack of domain knowledge, and error propagation (Schnoes et al. 2009).

Recently, the topic of misannotation in large-scale public protein sequence databases was brought to the attention of the scientific community (Pegg et al. 2006; Lundin et al. 2009; Schnoes et al. 2009; Nobre et al. 2016). For example, it was found that the rate of misannotation for enzyme superfamilies varies from 5% to >80% and is increasing, with overprediction of function being the largest problem (Schnoes et al. 2009). We examined the scale of misannotation in our set of fully-sequenced genomes of Proteobacteria using the well-conserved chaperonin GroEL. All sequences of this protein clustered in one cluster, and the various terms used to annotate this protein are shown in Table 7.

**Table 7.** Annotations of Proteins in the GroEL Cluster and Other Proteins Labelled GroEL.

| Annotation | # Sequences | Percent | Notes |
| --- | --- | --- | --- |
| **Cluster 3128:** |  |  |  |
| GroEL | 1934 | 67.39% |  |
| GroL | 410 | 14.29% |  |
| 60 kDa chaperonin | 338 | 11.78% |  |
| chaperonin Cpn60/TCP-1 | 76 | 2.65% |  |
| chaperone Hsp60 | 26 | 0.91% |  |
| chaperonin/chaperon/chaperone | 16 | 0.56% |  |
| heat shock protein | 8 | 0.28% |  |
| hypothetical | 7 | 0.24% |  |
| Stress protein H5 | 5 | 0.17% |  |
| thermosome | 2 | 0.07% |  |
| GroESL | 2 | 0.07% |  |
| mopA | 1 | 0.03% | Obsolete annotation |
| not yet annotated | 1 | 0.03% |  |
| **Other Clusters and Singletons:** |  |  |  |
| GroEL | 44 | 1.53% | Mislabeled |
| Total | 2870 | 100% |  |

Cluster 3128 contains 2,826 GroEL sequences, and 2,344 sequences (82%) contained the term “GroEL” or “GroL” in their annotation (Table 7). The remaining GroEL sequences had variable annotations including “not yet annotated” (CDS AHN71682.1, *Aggregatibacter actinomycetemcomitans* str. HK1651, Org. 1247). Genes for the GroEL protein are correctly annotated as either *groEL* or *groL.* As such, it is understandable that the protein has come to be widely annotated by either GroEL or GroL. However, SWISS-PROT indicates the correct annotation to be 60 kDa chaperonin while NCBI RefSeq refers to it as molecular chaperone GroEL. Thus, it is obvious that even between curated databases there is not a consensus on the correct annotation for a universally conserved protein. Lund reports that GroEL should be used exclusively for *Escherichia coli* (Lund 2009), and he recommends the use of Cpn60 on the basis of a nomenclature system proposed by Coates et al. (Coates et al. 1993). However, we found that GroEL was used predominantly regardless of species.

The third most commonly used term was 60 kDa chaperonin, but it was only used in 11.78% of the annotations. Other terms that were used included chaperonin; Cpn60/Tcp-1, the latter of which is a term used for eukaryotic molecular chaperones; various constructs of the word chaperone; heat shock protein which is a general term for describing the function of chaperones; and mopA which is an obsolete gene name for *groEL*, and in seven annotations it was not recognized as GroEL and annotated as hypothetical (Table 7).

Forty-four more sequences incorrectly annotated as “chaperonin GroEL” were found in other clusters or as singletons. Characteristics of these clusters and singletons indicate that none of them contain true GroEL proteins. Examples are proteins ABR79862.1, AEG96676.1, and AKH10420.1 belonging to *Klebsiella pneumoniae* subsp. Pneumoniae str. MGH 78578 (Org. 227), *Enterobacter aerogenes* str. KCTC 2190 (Org. 700), and *Salmonella enterica* subsp. Enterica (Org. 1845), respectively. All three proteins are annotated “chaperonin GroEL”; however, none of them are.

## Discussion

In this study we examined the distribution and the level of conservation among four essential proteins – GroEL, RpoB/RpoB’, and PolA. We found that GroEL is the most conserved protein in the set. All microorganisms have GroEL, and its sequences clustered in a single cluster indicating high conservation of biological function. Thus, for GroEL we accept our hypothesis that this protein is conserved in sequence and function across all Proteobacteria. However, we found that for the other three proteins in this study, proteins clustered in multiple clusters or were missing indicating that there was a range of sequence and structure that could perform the function of the particular protein. Therefore, for these three proteins, we reject our hypothesis.

Many of the instances in which we found alternative clusters for protein function or a missing sequence occurred in endosymbionts. Endosymbionts typically have reduced genomes and have eliminated genes that are not necessary for their obligate intracellular lifestyle (Wernegreen 2002; Tamas et al. 2008). Thus, it is not surprising that when we found unusual arrangements of genes or else missing genes, these occurred most often in endosymbionts. One common explanation for how endosymbionts can exist in the absence of genes and gene products is that they import necessary functions from their host. For the particular proteins in our study, we do not know how the endosymbionts fulfill the functions when the genes are missing.

There were a few instances of non-endosymbiont organisms that were missing either PolA or RpoB/RpoB’ and their respective genes. We suspect that the absence of the genes for these proteins was due to genome assembly issues. In the case of PolA, all Proteobacteria except for *M. haemolytica* str. USMARC_2286 (Org. 1073) contained the protein, while in the case of RpoB/RpoB’ it was more difficult to discern as the unusual species *(“Ca.* F. marinum” (Orgs. 1822, 1823)) did not have any close relatives with good synteny or sequence identity in the NCBI database.

Our study highlights difficulties that can be encountered when working with genomes on a large scale due to errors that can occur at two different points during the acquisition or curation of data. Clearly a sequence is only as good as technology allows, and as we have indicated several times, a number of the genes in our study contained frameshifts for which we cannot know were real or due to technical sequencing errors. We know that read errors in homopolymeric tracks are frequent with high throughput sequencing technologies and can result in frameshifts (Henson et al. 2012). We suspect many of the frameshifts we encountered are due to sequence errors because there were closely related sequences that contained intact genes. The second issue of annotation is a more open problem that includes both human and algorithm error. Examples of algorithm error are problems such as recognizing the correct open reading frame for a gene or the proper start site. Many researchers are now using automated programs to functionally annotate their genomes when they have very little knowledge of a given gene or gene family, and when faced with ambiguous information on which to make a decision about naming a gene product, they make a choice based on closest BLAST hits or association with a similar species; this can propagate errors (Schnoes et al. 2009). We’ve illustrated a number of these situations throughout our study.

The annotation errors we uncovered provide a cautionary tale to anyone working with genomes on a large scale. Big data scientists need to be able to rely on the data they are using to be correctly identified, and with the level of misannotation in public databases, genome data may not be reliable for some big data applications. For example, only 0.1% of the 224,442 clusters contained sequences from at least 90% of the 2,307 organisms in this study, approximately 225 clusters each representing a unique protein sequence. Given the results of our analyses, the actual percentage of clusters containing at least 90% of the organisms is certainly higher.

## Methods

Protein sequences of all completed Proteobacterial genomes and their associated plasmids were downloaded from NCBI GenBank ftp service (Benson et al. 2005) in the second half of February 2016. Every microorganism in this study was assigned an organism number that can be found in (Supplemental Table S2). 8.76 M protein sequences were aligned with a distributed memory software pGraph-Tascel (Daily 2016) and clustered with Grappolo (Lu et al. 2015).

### Alignment with pGraph-Tascel

The analysis pipeline for the pairwise alignment of 8.76 M sequences involved a number of automated steps carried out by the pGraph-Tascel software (Daily 2016). The steps include: 1) filtering out pairs of sequences that can be accurately predicted to produce poor alignment scores, 2) performing the alignments of the remaining sequences, and finally 3) filtering out the alignments that do not meet the cutoff criteria. The acceptance criteria for an alignment is that its length must be 80% of the longer sequence, 40% of the alignment must be exact matches, and the alignment must achieve at least 30% of the optimal score of the longer sequence aligned against itself. These acceptable alignments represent an edge in our output graph with the two sequences representing the vertexes of the edge.

There are two criteria for filtering out unwanted sequence pairs prior to alignment. The first criterion is a simple length filter. Since the eventual edge criterion depends on the length of the alignment relative to the longest sequence, we can eliminate alignment pairs when the lengths of two sequences differ more than the eventual edge criterion would eliminate. The second filter criterion eliminates sequence pairs if the sequences do not contain an exact-matching subsequence of a given minimum length. In practice, a cutoff length of seven worked well for protein sequences. A suffix array data structure is used to determine which sequence pairs meet the exact-match length criteria.

The remaining sequence pairs that are suitable for alignment can be aligned efficiently using standard parallel processing techniques since the alignments are naturally independent of each other. The performance is further improved by using the Parasail library (Daily et al. 2015). The Parasail library implements pairwise sequence alignment using special vector instruction sets of commodity CPUs, resulting in significant performance improvements of 10x and greater over non-vectorized implementations.

The pGraph-Tascel software orchestrates this entire filter-and-align pipeline. When processing large sets of sequences such as the 8.76 M presented here, the problem can be efficiently run on modest-sized compute clusters. In this case, the set of sequences is broken into smaller tiles, where each tile represents the entire filter-and-align pipeline. These tiles of work can then be independently solved by any available CPU in the compute cluster. Because there is a variable amount of work for any given set of sequences, any load-balancing issues are mitigated by a distributed-memory task counter, enumerating the tiles and handing off the next available task when an idle CPU requests the work (Daily 2016). The complete pGraph-Tascel suite with installation instructions is available at https://github.com/jeffdaily/parasail. The parasail_aligner module for sequence alignment is available at https://github.com/jeffdaily/parasail/blob/master/apps/README.md.

Once the sequences have been aligned and transformed into a homology graph using the criteria described previously, the graph is processed by the Grappolo community detection software (Lu et al. 2015).

### Clustering with Grappolo

The output of pGraph-Tascel is an undirected graph *G*(*V, E*), where *V* is the set of input protein sequences and *E* is the set of edges (*v*__*i*__, *V*_*j*_) such that the sequences *v*__*i*__ and *V*_*j*_ are similar based on the specified criteria. The graph *G* serves as input to Grappolo - a dense subgraph detection algorithm that forms clusters using sequence similarity measure (Lu et al. 2015).

Grappolo implements parallelization of the Louvain heuristic (Blondel et al. 2008) for community detection in large-scale graphs (Lu et al. 2015). The algorithm finds communities by optimizing the modularity metric (Newman and Girvan 2004). Intuitively, modularity measures the fraction of the within-community edges minus the expected value of random edges between the vertices in a network with the same community divisions (Newman and Girvan 2004). Although, modularity is not an ideal measure, it seems to work well in practice.

In our application, a “community” is a set of closely related protein sequences. Thus, Grappolo clusters protein sequences using similarity measure computed by pGraph-Tascel. In our application we used the alignment length statistic as the edge weight. Grappolo has been shown to produce clusters of high modularity (Lu et al. 2015). The clusters created by Grappolo contain proteins that are closely related in sequence as well as in function. Grappolo software is available at https://github.com/luhowardmark/GrappoloTK.

### Cluster post-processing

Cluster and sequence information were stored in a cluster text file. For each protein in the study, the cluster text file was queried using one or more regular expressions characteristic of the protein annotations (Supplemental File S1). This procedure identified clusters of potential interest that were subsequently extracted. Because protein annotations are not a very reliable source for determining protein function, the extracted clusters were manually inspected for relevance, and false positives were removed, i.e., clusters containing sequences that were misannotated as sequences of interest but were not. The remaining clusters were analyzed.

## Acknowledgments

The authors wish to thank all of the dedicated researchers who contributed completed genomes to the NCBI database. While the rate of genome data is growing at a phenomenal rate, the rate of completed genomes is growing much more slowly because this work is that much more difficult. While we have reported on issues with annotation, this is not meant as a slight to any researcher, but just a realization of the difficulties of this task. This work received support from the National Science Foundation under the Advances in Biological Informatics program, Award DBI 1262664.

## References

Altschul, SF, Gish, W, Miller, W, Myers, EW, Lipman, DJ. 1990. Basic local alignment search tool. J Molec Biol 215: 403–410.

Apetri, AC, Horwich, AL. 2008. Chaperonin chamber accelerates protein folding through passive action of preventing aggregation. Proc Natl Acad Sci U S A 105: 17351–17355.

Benson, DA, Karsch-Mizrachi I, Lipman, DJ, Ostell, J, Wheeler, DL. 2005. GenBank. Nucl Acids Res 33: D34–D38.

Berg, JM, Tymoczko, JL, Stryer, L. 2002. Transcription Is Catalyzed by RNA Polymerase. W. H. Freeman, New York.

Bernad, A, Blanco, L, Lázaro, JM, Martin, G, Salas, M. 1989. A conserved 3’→5’ exonuclease active site in prokaryotic and eukaryotic DNA polymerases. Cell 59: 219–228.

Bhutani, N, Udgaonkar, JB. 2002. Chaperonins as protein-folding machines. Curr Sci: 1337–1351.

Blondel, VD, Guillaume, J-L, Lambiotte, R, Lefebvre, E. 2008. Fast unfolding of communities in large networks. J Stat Mech 2008: P10008.

Camps, M, Loeb, LA. 2004. When pol I goes into high gear: processive DNA synthesis by pol I in the cell. Cell Cycle 3: 114–116.

Chaparro, PJP, McCulloch, JA, Cerdeira, LT, Al-Dilaimi A, de Sá LLC, de Oliveira R, Tauch, A, de Carvalho Azevedo, VA, Schneider, MPC, da Silva ALdC. 2011. Whole genome sequencing of environmental Vibrio cholerae O1 from 10 nanograms of DNA using short reads. J Microbiol Methods 87: 208–212.

Chapman, E, Farr, GW, Usaite, R, Furtak, K, Fenton, WA, Chaudhuri, TK, Hondorp, ER, Matthews, RG, Wolf, SG, Yates, JR. 2006. Global aggregation of newly translated proteins in an Escherichia coli strain deficient of the chaperonin GroEL. Proc Natl Acad Sci U S A 103: 15800–15805.

Coates, ARM, Shinnick, TM, Ellis, RJ. 1993. Chaperonin nomenclature. Mol Microbiol 8: 787–787.

Cooper, GM. 2000. The Cell: A Molecular Approach. In The Cell: A Molecular Approach. Sinauer Associates, Sunderland, MA.

Daily, J. 2016. Parasail: SIMD C library for global, semi-global, and local pairwise sequence alignments. BMC Bioinformatics 17: 81.

Daily, J, Kalyanaraman, A, Krishnamoorthy, S, Vishnu, A. 2015. A work stealing based approach for enabling scalable optimal sequence homology detection. J Parallel Distrib Comput 79: 132–142.

Fenton, WA, Horwich, AL. 2003. Chaperonin-mediated protein folding: fate of substrate polypeptide. Q Rev Biophys 36: 229–256.

Friedberg, EC, Walker, GC, Siede, W, Wood, RD. 2005. DNA repair and mutagenesis. American Society for Microbiology Press.

Gil, R, Silva, FJ, Peretó, J, Moya, A. 2004. Determination of the core of a minimal bacterial gene set. Microbiol Mol Biol Rev 68: 518–537.

Helmann, JD, Chamberlin, MJ. 1988. Structure and function of bacterial sigma factors. Annu Rev Biochem 57: 839–872.

Henson, J, Tischler, G, Ning, Z. 2012. Next-generation sequencing and large genome assemblies. Pharmacogenomics J 13: 901–915.

Hübscher, U, Spadari, S, Villani, G, Maga, G. 2010. DNA Polymerases in the Three Kingdoms of Life:Bacteria, Archaea and Eukaryotes. In DNA Polymerases, doi:10.1142/9789814299176_0002, pp. 59–83. WORLD SCIENTIFIC.

Hutchison, CA, 3rd, Chuang, RY, Noskov, VN, Assad-Garcia, N, Deerinck, TJ, Ellisman, MH, Gill, J, Kannan, K, Karas, BJ, Ma, L et al. 2016. Design and synthesis of a minimal bacterial genome. Science 351: aad6253.

Kogoma, T. 1997. Stable DNA replication: interplay between DNA replication, homologous recombination, and transcription. Microbiol Mol Biol Rev 61: 212–238.

Koonin, EV. 2000. How many genes can make a cell: the minimal-gene-set concept. Annu Rev Genomics Hum Genet 1: 99–116.

Koonin, EV. 2003. Comparative genomics, minimal gene-sets and the last universal common ancestor. Nat Rev Microbiol 1: 127.

Lin, Z, Madan, D, Rye, HS. 2008. GroEL stimulates protein folding through forced unfolding. Nat Struct Mol Biol 15: 303.

Lu, H, Halappanavar, M, Kalyanaraman, A. 2015. Parallel heuristics for scalable community detection. Parallel Comput 47: 19–37.

Lund, PA. 2009. Multiple chaperonins in bacteria-why so many? FEMS Microbiol Rev 33: 785–800.

Lundin, D, Torrents, E, Poole, AM, Sjöberg B-M. 2009. RNRdb, a curated database of the universal enzyme family ribonucleotide reductase, reveals a high level of misannotation in sequences deposited to Genbank. BMC Genomics 10: 589.

Newman, MEJ, Girvan, M. 2004. Finding and evaluating community structure in networks. Phys Rev E 69: 026113.

Nobre, T, Campos, MD, Lucic-Mercy, E, Arnholdt-Schmitt B. 2016. Misannotation awareness: a tale of two gene-groups. Front Plant Sci 7: 868.

Pagnier, I, Yutin, N, Croce, O, Makarova, KS, Wolf, YI, Benamar, S, Raoult, D, Koonin, EV, La Scola B. 2015. Babela massiliensis, a representative of a widespread bacterial phylum with unusual adaptations to parasitism in amoebae. Biology Direct 10: 13.

Pegg SC-H, Brown, SD, Ojha, S, Seffernick, J, Meng, EC, Morris, JH, Chang, PJ, Huang, CC, Ferrin, TE, Babbitt, PC. 2006. Leveraging enzyme structure function relationships for functional inference and experimental design: the structure- function linkage database. Biochemistry 45: 2545–2555.

Schnoes, AM, Brown, SD, Dodevski, I, Babbitt, PC. 2009. Annotation error in public databases: misannotation of molecular function in enzyme superfamilies. PLoS Comput Biol 5: e1000605.

Shigenobu, S, Watanabe, H, Hattori, M, Sakaki, Y, Ishikawa, H. 2000. Genome sequence of the endocellular bacterial symbiont of aphids Buchnera sp. APS. Nature 407: 81.

Smalley, DJ, Whiteley, M, Conway, T. 2003. In search of the minimal Escherichia coli genome. Trends Microbiol 11: 6–8.

Sutton, MD, Walker, GC. 2001. Managing DNA polymerases: coordinating DNA replication, DNA repair, and DNA recombination. Proc Natl Acad Sci U S A 98: 8342–8349.

Sydow, JF, Cramer, P. 2009. RNA polymerase fidelity and transcriptional proofreading. Curr Opin Struct Biol 19: 732–739.

Tamas, I, Klasson, L, Canbäck, B, Näslund, AK, Eriksson A-S, Wernegreen, JJ, Sandström, JP, Moran, NA, Andersson, SGE. 2002. 50 million years of genomic stasis in endosymbiotic bacteria. Science 296: 2376–2379.

Tamas, I, Wernegreen, JJ, Nystedt, B, Kauppinen, SN, Darby, AC, Gomez-Valero L, Lundin, D, Poole, AM, Andersson, SGE. 2008. Endosymbiont gene functions impaired and rescued by polymerase infidelity at poly (A) tracts. Proc Natl Acad Sci U S A 105: 14934–14939.

van Ham RCHJ, Kamerbeek, J, Palacios, C, Rausell, C, Abascal, F, Bastolla, U, Fernández, JM, Jiménez, L, Postigo, M, Silva, FJ. 2003. Reductive genome evolution in Buchnera aphidicola. Proc Natl Acad Sci U S A 100: 581–586.

Wernegreen, JJ. 2002. Genome evolution in bacterial endosymbionts of insects. Nat Rev Genet 3: 850.

